# Investigation of electrochromic band-shifts in the Soret region induced by the formation of Tyr_D_^•^, Tyr_Z_^•^, and Q_A_^•-^ in Photosystem II

**DOI:** 10.1101/2024.11.21.624785

**Authors:** Alain Boussac, Takumi Noguchi, A. William Rutherford, Julien Sellés, Miwa Sugiura, Stefania Viola

**Affiliations:** Institut de Biologie Intégrative de la Cellule, UMR9198, CEA Saclay, 91191 Gif-Sur-Yvette, France; Department of Physics, Graduate School of Science, Nagoya University, Furo-cho, Chikusa-ku, Nagoya 464-8602, Japan.; Department of Life Sciences, Imperial College, SW7 2AZ London, UK.; Institut de Biologie Physico-Chimique, UMR CNRS 7141 and Sorbonne Université, 13 rue Pierre et Marie Curie, 75005 Paris, France.; Proteo-Science Research Center, and Department of Chemistry, Graduate School of Science and Technology, Ehime University, Bunkyo-cho, Matsuyama, Ehime 790-8577, Japan.; Institute of Biosciences and Biotechnologies of Aix-Marseille – UMR7265, Saint-Paul-Lez-Durance, France.

## Abstract

The effects of Tyr_D_^•^, Tyr_Z_^•^, and Q_A_^•-^ radical formation on the absorption spectrum in the Soret region were studied in Mn-depleted Photosystem II at pH 8.6 (in order to be in the Tyr_D_ state after dark adaptation). Flash-induced difference spectra were recorded in several PSII samples from: *i*) *Thermosynechococcus vestitus* (formerly *T. elongatus*), *ii*) *Synechocystis sp.* PCC 6803, *iii*) *Chroococcidiopsis thermalis* PCC 7203 grown under far-red light, and *iv*) *Acaryochloris marina*. In the case of *T. vestitus*, mutants D1/H198Q, D1/T1789H, D2/I178H, and D2/Y160F, with PsbA1/Q130 instead of PsbA3/E130, were also studied for possible contributions from P_D1_, Chl_D1_, Chl_D2_, and Phe_D1_, respectively. For a possible contribution from P_D2_, the D2/H197A mutant was studied in *S. 6803*. While P_D1_ is clearly the species whose spectrum is blue-shifted by ∼3nm in the presence of Q_A_^•-^, as has already been well documented in the literature, the species whose spectra shift upon the formation of Tyr_D_^•^ and Tyr_Z_^•^ remain to be clearly identified, as they appear different from P_D1_, P_D2_, Phe_D1_, Chl_D1_, and Chl_D2_, as concluded by the lack of different light-induced difference spectra in the mutants listed above. Although we cannot rule out a weak effect, considering the accuracy of the experiments, it is proposed that other pigments, such as antenna Chl and/or Car, near the reaction center are involved. Additionally, it is shown that: *i*) there is no proton release into the bulk upon the oxidation of Tyr_D_ at pH 8.6, and *ii*) the rearrangement of the electrostatic environment of the pigments involved in the light-induced difference spectra in the samples studied, upon the formation of Tyr_D_^•^, Tyr_Z_^•^, and Q_A_^•-^, likely occurs differently from both a kinetic and structural perspective.

## Introduction

Oxygenic photosynthesis in cyanobacteria, algae, and higher plants converts solar energy into the chemical bonds of sugars and dioxygen. Photosystem II (PSII) begins this process by splitting water to obtain electrons in the form of reduced quinones, while generating a proton gradient and releasing O . The mature PSII binds 35 chlorophyll *a* (Chl- *a*), 2 pheophytins (Phe-*a*), 1 membrane b-type cytochrome, 1 extrinsic c-type cytochrome (in cyanobacteria and red algae), 1 non-heme iron, 2 plastoquinones (Q_A_ and Q_B_), the Mn CaO cluster, 2 Cl, 12 carotenoids, and 25 lipids [1,2]. In the cyanobacterium *Synechocystis sp.* PCC 6803, a fourth extrinsic subunit, PsbQ, has also been found in addition to PsbV, PsbO, and PsbU [3].

Among the 35 Chl-a molecules, 31 are antenna Chl’s. When one is excited, the excitation energy is transferred to other chlorophylls until it reaches the key pigments in the photochemical reaction center [3]: 4 Chl-*a* molecules, P_D1_, P_D2_, Chl_D1_, Chl_D2_, and 2 Phe-*a* molecules, Phe_D1_ and Phe_D2_. A few picoseconds after the formation of the excited *Chl_D1_, charge separation occurs, ultimately resulting in the formation of the Chl_D1_^•+^Phe_D1_^•-^ and then P_D1_^•+^Phe_D1_^•-^ radical pair states [5,6]. The formation of the Chl_D1_^•+^Phe_D1_^•-^ radical pair was recently defined as a fast pathway (short-range charge separation) in contrast with a slow pathway where P_D1_P_D2_ acts as the initial donor (long-range charge separation), which would result directly in the formation of the P_D1_^•+^Phe_D1_^•-^ radical pair [7].

After this charge separation, P_D1_^•+^ oxidizes Tyr_Z_, the Tyr161 of the D1 polypeptide, which is then reduced by the Mn CaO cluster. The electron on Phe_D1_^•-^ is then transferred to Q_A_, the primary quinone electron acceptor, and then to Q_B_, the secondary quinone electron acceptor. While Q_A_ can only be singly reduced under normal conditions, Q_B_ accepts two electrons and two protons before leaving its binding site and being replaced by an oxidized Q_B_ molecule from the membrane plastoquinone pool. For a non-exhaustive list of recent reviews on PSII function, see for example [8–12]. The Mn CaO cluster, oxidized by the Tyr_Z_^•^ radical formed after each charge separation, cycles through five redox states denoted S_n_, where n represents the number of stored oxidizing equivalents. When the S_4_-state is formed, two water molecules bound to the cluster are oxidized, O is released, and the S_0_-state is reformed [13,14].

Formation of the Q_A_^•-^ species is known to induce spectral changes in Phe_D1_ in the Qx region around 550 nm [15–17], and in P_D1_ in the Qy region between 650 and 700 nm [18,19]. These electrochromic responses have been extensively studied both experimentally and theoretically For example, it has been observed that in the presence of Q_A_^•-^, the blue shift of the Qy band of Chl_D1_, typically seen at ∼680 nm, was replaced by one at ∼727 nm in far read light PSII (FRL-PSII), indicating that the Chl-*a* acting as Chl_D1_ was replaced by a FRL chlorophyll (likely a Chl-*d)* [20]. The Soret band of P_Dļ_ is also sensitive to the formation of Q_A_^•-^ and to the charge in and around the Mn CaO cluster [21,22]. That said, spectral changes in the blue region are far less well-studied than the Qx and Qy bands, although see [18] for further discussion. For this reason, we report here an initial characterization of the spectral changes between 400 and 460 nm induced by the formation of the Tyr_D_^•^, Tyr_Z_^•^, and Q_A_^•-^ radicals. In order to achieve the highest proportion of centers with a reduced Tyr_D_ after dark adaptation, and to avoid contributions from the Mn CaO cluster, the experiments were performed at pH 8.6 in several Mn-depleted PSII preparations [23].

## Materials and Methods

The PSII samples used were: *i*) the PsbA3-PSII [24], the PsbA3/H198Q mutant [25], the PsbA3/T179H mutant [26], the Tyr_D_-less mutant (with D1 = PsbA1) [27], and the D2/I178H mutant (with D1 = PsbA3) [28], all purified from *Thermosynechococcus vestitus*; *ii*) the FRL-PSII from *Chroococcidiopsis thermalis* [20,29], with 4 Chl-*f* and 1 Chl-*d*; *iii*) the wild-type and D2/H197A mutant from *Synechocystis* 6803 [30]; and *iv*) PSII from *Acaryochloris marina* [29], with 34 Chl-*d* and 1 Chl-*a* at the P_D1_ position [31].

The samples were Mn-depleted by adding approximately 10 mM NH OH (from a 1 M stock at pH 6.5) and 1 mM EDTA. The hydroxylamine was then removed by washing the PSII samples through cycles of dilution in 1 M betaine, 15 mM CaCl, 15 mM MgCl, 100 mM Tris, 100 µM EDTA, pH 8.6 (adjusted with HCl), followed by concentration using Amicon Ultra-15 centrifugal filter units (cut-off 100 kDa) until the estimated residual NH OH concentration was lower than 0.1 µM in the concentrated PSII samples before the final dilution. In the final dilution step, the PSII samples were suspended in 1 M betaine, 15 mM CaCl, 15 mM MgCl, 100 mM Tris, pH 8.6. For the experiment reported in Fig. 8, the Tris buffer was replaced with Mes buffer at pH 6.5.

For the measurements with the dye thymol blue, the Mn-depleted Tyr_D_-less PSII at pH 8.6 was washed through cycles of dilution in 100 mM NaCl, 15 mM CaCl, and 15 mM MgCl, followed by concentration using Amicon Ultra-15 centrifugal filter units (cut-off 100 kDa), until the estimated residual Tris concentration was lower than 1 µM in the concentrated PSII samples before the final dilution. After dark adaptation for 3 to 4 hours at room temperature, 150 µM thymol blue and 100 µM PPBQ (in dimethyl sulfoxide) were added. During the measurement, the sample was continuously bubbled with ALPHAGAZ 2 Argon from Air Liquide, in which the CO contamination is < 0.1 ppm. This bubbling was performed to minimize drift in the baseline (*i.e.*, in the pH) due to the solubilization of atmospheric CO .

Time-resolved absorption changes were measured with a lab-built spectrophotometer [32], modified as described in detail in [33]. The chlorophyll concentration of the samples was approximately 25 µg of Chl mL^1^.

## Results and Discussion

Panel A in Fig. 1 shows the ΔI/I recorded at 432 nm in Mn-depleted PsbA3-PSII at pH 8.6, illuminated by 10 saturating flashes (spaced 2 s apart) in the presence of the electron acceptor PPBQ. The sample was dark-adapted for 3 to 4 hours at room temperature before the flash illumination. Panel B shows a zoom of the data recorded after the first 4 flashes. In this experiment, the first weak, non-actinic detector flash was given 200 µs after each actinic flash. The second weak, non-actinic detector flash was given 10 ms after each actinic flash. Then, 20 weak, non-actinic detector flashes (spaced 100 ms apart) were given after each of the 10 actinic flashes.

**Figure 1:**
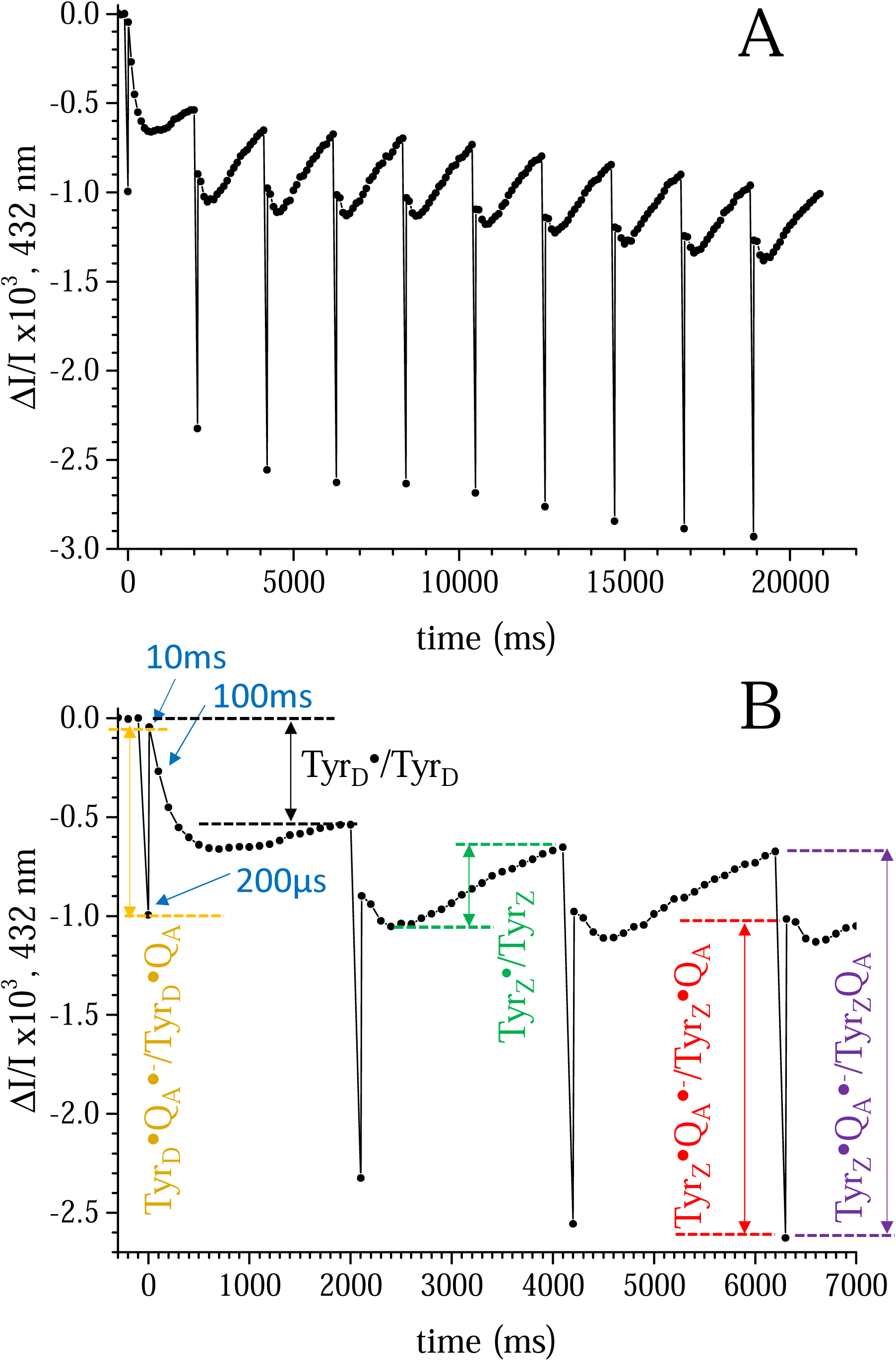
Panel A shows the ΔI/I changes recorded at 432 nm in Mn-depleted PsbA3-PSII from *T. vestitus* at pH 8.6. The sample was dark-adapted for 3 to 4 hours to allow Tyr_D_ to be almost fully reduced. After the dark adaptation, 100 µM of PPBQ, dissolved in dimethyl sulfoxide, were added. Flash illumination was performed with 10 saturating flashes (spaced 2 s apart). The first weak detector laser flash was given 200 µs after each of the saturating actinic laser flashes. The second detector flash was given 10 ms after each of the actinic flashes. Then, 20 detector flashes (spaced 100 ms apart) were given after each of the 10 actinic flashes. The optical density (OD) of the sample was 1.75 at 673 nm (∼25 µg Chl/ml). The pathlength of the cuvette was 2.5 mm. Panel B shows the same data as in Panel A, with an expanded time scale for the first 4 actinic flashes. The vertical black arrow, 2 s after the first flash, corresponds to the Δ(ΔI/I) mainly originating from the formation of Tyr_D_^•^ alone, since Q_A_^•-^ and Tyr_Z_^•^ (formed in centers with Tyr_D_^•^ present) are no longer present 2 s after the flash. After the 4th flash, when Tyr_D_^•^ has been formed in the majority of centers, the flash illumination induces the formation of the Tyr_Z_^•^Q_A_^•-^ state. The vertical red arrow shows the Δ(ΔI/I) between 200 µs and 10 ms and corresponds to the Tyr_Z_^•^Q_A_^•-^-*minus* -Tyr_Z_^•^Q_A_ difference. The vertical violet arrow shows the Δ(ΔI/I) between 200 µs and 2 s and corresponds to the Tyr_Z_^•^Q_A_^•-^-*minus*-Tyr_Z_Q_A_ difference. The vertical green arrow shows the Δ(ΔI/I) between 400 ms and 2 s and corresponds to the Tyr_Z_^•^-minus-Tyr_Z_ difference. The vertical orange arrow shows the Δ(ΔI/I) between 200 µs and 10 ms after the 1st actinic flash and corresponds to the Q_A_^•-^ *-minus*-Q_A_^•^ difference (in the presence of Tyr_D_^•^). The Δ(ΔI/I) between 10 ms and 400 ms will be analyzed in Fig. 8.

At pH 8.6, and after a long dark-adaptation, Tyr_D_ is reduced in the great majority of the centers. Under these conditions, the first actinic flash oxidizes either Tyr_D_ or Tyr_Z_ in almost equal proportions [23]. The non-decaying ΔI/I recorded 2 s after the first flash (black vertical arrow in Panel B of Fig. 1) is a consequence of Tyr_D_^•^ formation alone, since 2 s after the flash, Tyr_Z_^•^ and QQ_A_^•-^ are no longer present. Always after the first flash, the yellow vertical arrow (the 200 µs-*minus* -10 ms difference) corresponds to a mixture of the Tyr_D_^•^Q_A_^•-^-*minus* -Tyr_D_^•^Q_A_ and Tyr_Z_^•^Q_A_^•-^-*minus*-Tyr_Z_^•^Q_A_ differences. In other words, it is a mix of the Q_A_^•-^- *minus*-Q_A_ difference in the presence of either Tyr_D_^•^ or Tyr_Z_^•^. After the 4th flash, when Tyr_D_^•^ is formed in virtually all centers, the flash illumination only induces the formation of the Tyr_Z_^•^Q_A_^•-^ state, and the slow decay in the second (s) time range corresponds to the decay of Tyr_Z_^•^. Consequently, the red vertical arrow (the 200 µs-*minus*-10 ms difference) corresponds to the Tyr_Z_^•^Q_A_^•-^-*minus*-Tyr_Z_^•^Q_A_ difference, in other words, the Q_A_^•-^-*minus*-Q_A_ difference in the presence of Tyr_Z_^•^ and Tyr_D_^•^. This differs from the first flash, where the Q_A_^•-^-*minus*-Q_A_ difference was measured in the presence of only one of the two tyrosyl radicals, either Tyr_Z_^•^ or Tyr_D_^•^. Always after the 4th flash, the violet vertical arrow in Panel B of Fig. 1 is the ΔI/I 200 µs after flash n *minus* 2 s after flash n-1. It corresponds to the Tyr_Z_^•^Q_A_^•-^-*minus*-Tyr_Z_Q_A_ difference. Finally, the green vertical arrow (300 ms-*minus*-2 s) in Panel B of Fig. 1 corresponds to Tyr_Z_^•^-*minus*-Tyr_Z_ difference alone (in the presence of Tyr_D_^•^).

The lower ΔI/I amplitude observed 200 µs after the first flash, compared with subsequent flashes, was most likely due to centers where non-heme iron was oxidized prior to flash illumination. Indeed, in these centers, the electron transfer from Q_A_^•-^ to Fe^3+^ is completed well before 200 µs [34,35].

The experiment presented in Fig. 1 was repeated for wavelengths ranging from 401 nm to 457 nm. To improve the signal-to-noise ratio, except for the data corresponding to the first flash, the spectra were averaged from the 6th to the 10th flashes. Fig. 2 shows the five spectra corresponding to the electrochromic responses defined in Panel B of Fig. 1, using the same color code.

**Figure 2:**
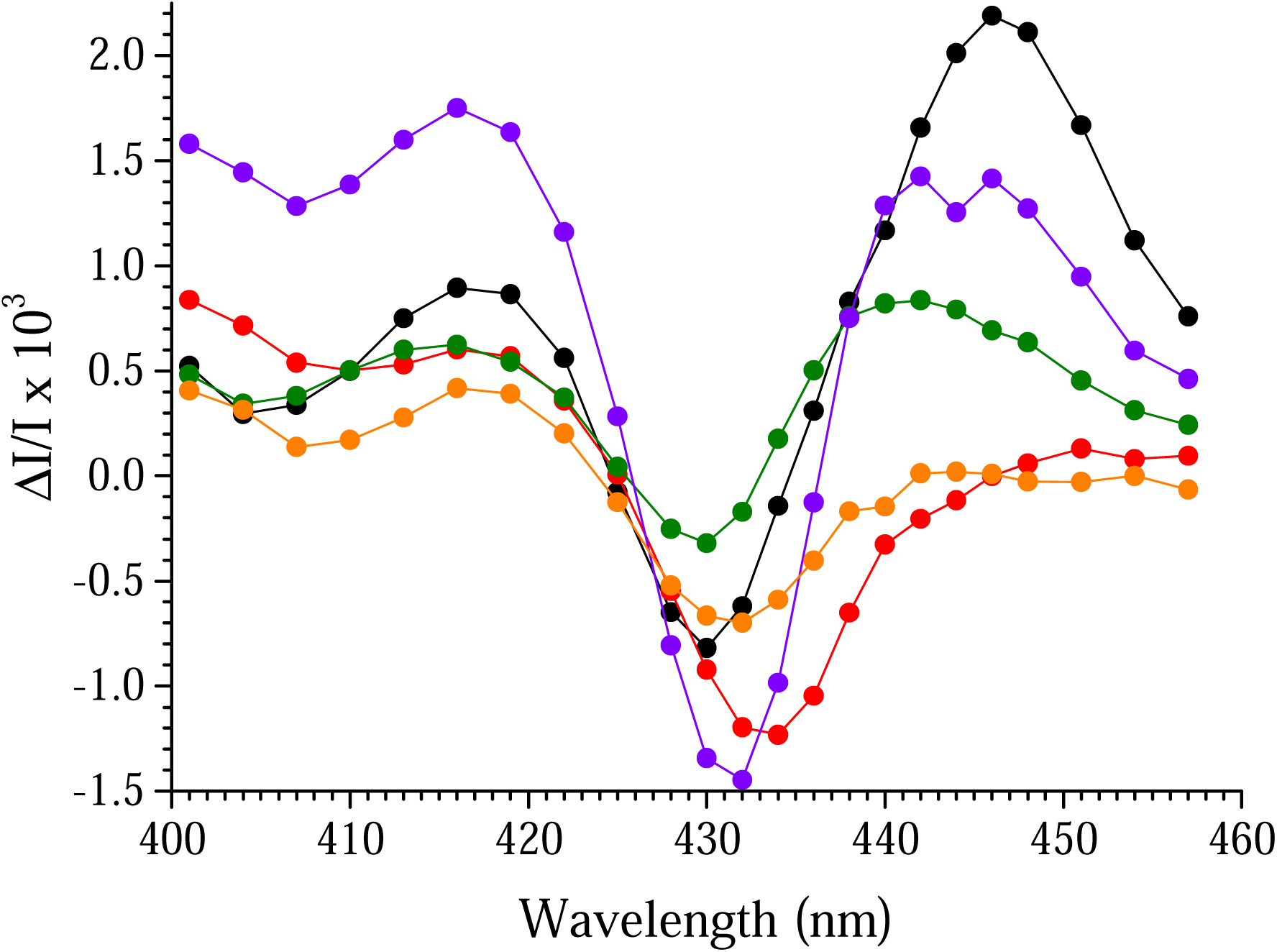
The experiment conducted at 432 nm with PsbA3-PSII from *T. vestitus*, as reported in Panel A of Fig. 1, was repeated for wavelengths between 401 nm and 457 nm. The different Δ(ΔI/I) values, defined in Panel B of Fig. 1, were plotted using the color code established in Panel B of Fig. 1. Except for the spectra corresponding to the first flash, the Δ(ΔI/I) values were averaged from the 6th to the 10th flash.

The spectrum in red, corresponding to the Q_A_^•-^-*minus*-Q_A_ difference in the presence of both Tyr_Z_^•^ and Tyr_D_^•^, may raise some questions. Indeed, it appears somewhat similar to the difference spectrum [P_D1_P_D2_]^•+^Q_A_^•-^-*minus*-[P_D1_P_D2_]Q_A_ recorded 20 ns after a flash [36]; see also [18]. The amplitude of the [P_D1_P_D2_]^•+^Q_A_^•-^ spectrum at 432 nm, recorded 20 ns after the flash under the same setup and with a similar sample, was ∼0.016 ΔI/I [36], approximately 11 times larger than the red spectrum observed here. The presence of ∼10% [P_D1_P_D2_]^•+^ at 200 µs after the flash could thus account for the ΔI/I signal recorded at 432 nm. However, attributing the species responsible for the slow decay in the 432-446 nm range is challenging [36]. See, however, the data at 820 nm [37]. Specifically, the measured decay corresponds to the [P_D1_P_D2_]^•+^Q_A_^•-^ spectrum, which necessarily also includes spectral contributions arising from Q_A_^•-^ formation. Most likely, these contributions are responsible for the phases observed in the hundreds of microseconds time range, which correspond to the electron transfer from Q_A_^•-^ to Q_B_. (There is no transfer to Q_B_^•-^, which is not present due to the addition of PPBQ, *e.g.*, [38].)

Additionally, the red spectrum in Fig. 2 closely resembles, both in shape and amplitude, the electrochromic shift induced by Q_A_^•-^ formation, attributed to a change in P_D1_ absorption under conditions where P_680_^+^ is excluded [22].

The Q_A_^•-^-*minus*-Q_A_ difference spectrum after the first flash (orange spectrum) shows a 2 nm blue shift compared to the red spectrum, from ∼434 nm to ∼432 nm. This indicates that the presence of both tyrosyl radicals shifts the Q_A_^•-^-*minus* -Q_A_ difference spectrum to the red compared to the condition with Tyr_D_^•^ alone. As explained earlier, after the first flash, there is a mixture of two difference spectra because Tyr_Z_^•^ and Tyr_D_^•^ are formed in approximately equal proportions [23]. This implies that the difference between the first flash spectrum and the average spectrum from the 6th to the 10th flashes would have been larger if Tyr_D_^•^Q_A_^•-^ had been formed exclusively after the first flash.

It is also noteworthy that the [P_D1_P_D2_]^•+^ spectrum measured 20 ns after a flash shows a dip at the same wavelength (∼432 nm) after both the first and subsequent flashes [36]. This provides additional evidence against attributing the orange and red spectra to [P_Dι_P_D2_]^•+^.

While the pigment whose absorption is modified by Q_A_^•-^ formation has been clearly identified as P_Di_ (*e.g.*, [21,22]), the pigment(s) affected by the formation of Tyr_Z_^•^ and Tyr_D_^•^ remain unidentified. This is the focus of the experiments reported below, which use PSII mutants and PSII from different species.

Panel A of Fig. 3 shows the ΔI/I signal recorded at 432 nm in Mn-depleted, Tyr_D_-less PSII [27] at pH 8.6, illuminated with 10 saturating flashes (spaced 2 seconds apart) in the presence of the electron acceptor PPBQ. Despite the absence of Tyr_D_ in this PSII, the sample was dark-adapted for 3–4 hours at room temperature before flash illumination.

**Figure 3:**
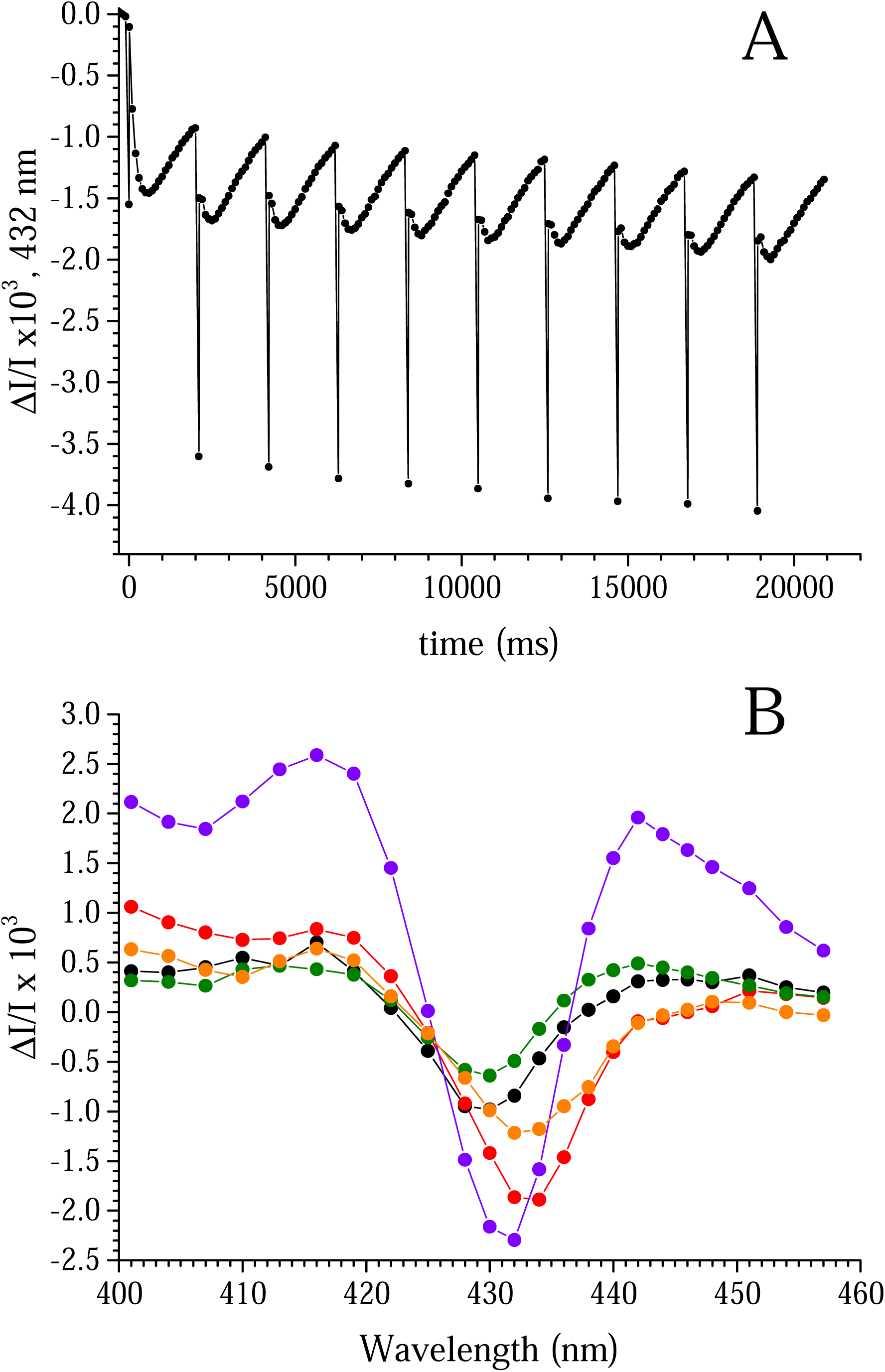
Panel A shows the ΔI/I changes recorded at 432 nm in Mn-depleted, Tyr_D_-less PSII from *T. vestitus* at pH 8.6, using the same protocol as described in Panel A of Fig. 1. Panel B shows the ΔI/I measured as in Panel A, for wavelengths between 401 nm and 457 nm, and plotted using the color code defined in Panel B of Fig. 1. Except for the spectra corresponding to the first flash, the Δ(ΔI/I) values were averaged from the 6th to the 10th flash.

Panel B of Fig. 3 displays the spectra constructed as described in Panel B of Fig. 1. Due to the absence of Tyr_D_, Tyr_Z_^•^ was formed after the first flash. The black spectrum recorded 2 seconds after the first flash is therefore attributed to the Tyr_Z_^•^- *minus*-Tyr_Z_ difference spectrum (green spectrum).

Two additional observations can be made: *i*) The Q_A_^•-^-*minus*-Q_A_ difference spectrum after the first flash (orange spectrum) is identical to the Q_A_^•-^-*minus*-Q_A_ difference spectrum recorded after subsequent flashes. *ii*) The trough of this spectrum in the presence of Tyr_Z_^•^ alone is located at ∼433 nm, which is intermediate between the Q_A_^•-^-*minus*-Q_A_ difference spectra observed in the presence of either Tyr_D_^•^ alone or both Tyr_D_^•^ and Tyr_Z_^•^.

This experiment demonstrates that the Q_A_^•-^-*minus*-Q_A_ difference spectrum shows slight variations across three conditions: with Tyr_D_^•^ alone, Tyr_Z_^•^ alone, or both Tyr_D_^•^ and Tyr_Z_^•^.

In Mn-depleted PSII at pH 6.5, the oxidation of both Tyr_Z_ and Tyr_D_ is coupled to proton release into the bulk [39,40]. In [39], it was proposed that the proton released upon Tyr_D_^•^ formation is directly released from the protein rather than being transferred to the neighboring D2-His198, unlike the process during Tyr_Z_^•^ formation, where the proton is transferred to the hydrogen-bonded D1-His190.

At pH 8.6, a pH value at which Tyr_D_^•^ is unstable in the dark, no information is currently available about the proton release following Tyr_D_ oxidation. Fig. 4 presents an experiment that tracks proton release into the bulk at pH 8.6, using the dye thymol blue, after a series of flashes applied to dark-adapted Mn-depleted PsbA3-PSII. At 580 nm, medium acidification results in a decrease in thymol blue absorption.

**Figure 4:**
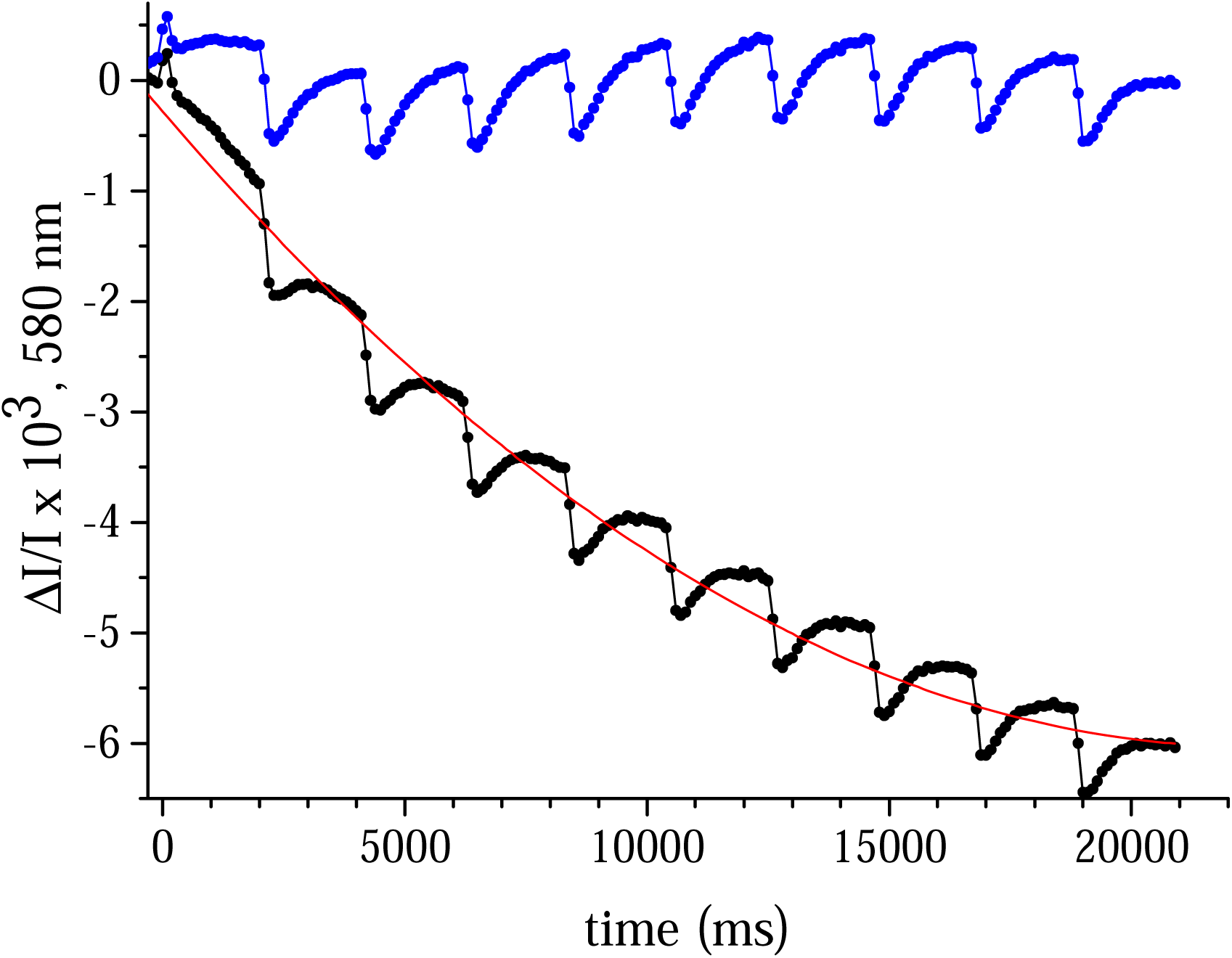
ΔI/I changes recorded at 580 nm in the presence of thymol blue (black data points) with dark-adapted Tyr_D_-less PSII from *T. vestitus* at pH 8.6. The first detector flash was given 20 µs after each actinic flash. Then, 20 detector flashes (spaced 100 ms apart) were given after each of the 10 actinic flashes. The continuous red curve corresponds to a second-degree polynomial fit. The blue trace represents the black trace *minu*s the red curve. This procedure was used to subtract as much drift as possible from the black trace.

The sample was illuminated with ten flashes, spaced 2 seconds apart, and the ΔI/I was measured every 100 ms (the first detection occurred 20 µs after the actinic flashes). The black trace represents the raw data. The red curve corresponds to a polynomial fit, and the blue trace is obtained by subtracting the polynomial fit from the black trace. This subtraction was performed to minimize drift in the black trace.

Despite bubbling the PSII suspension with pure argon gas, the drift likely results from the solubilization of trace amounts of atmospheric CO_2_.

The first conclusion from the experiment shown in Fig. 4 is that Tyr_D_ oxidation is not coupled to proton release into the bulk, as evidenced by the absence of a color change in thymol blue after the first flash. This contrasts with the oxidation of Tyr_Z_ during subsequent flashes, where a proton was released with kinetics not resolved in this experiment. The decay of Tyr_Z_^•^ was then coupled to proton uptake by PSII, with kinetics similar to those observed at 432 nm in Fig. 1A. To our knowledge, this is the first measurement of proton release and uptake at pH 8.6 associated with the oxidation of Tyr_D_ and subsequently Tyr_Z_.

The second conclusion is that both Tyr_Z_ and Tyr_D_ oxidations have an electrochromic effect on one or more pigments nevertheless the observation that in the case of Tyr_Z_ oxidation, the proton is released into the medium, whereas in the case of Tyr_D_, no proton release occurs.

In the following sections, we present comparisons of the different spectra defined in Panel B of Fig. 1, obtained from Mn-depleted PSII in mutants and PSII with different pigment compositions. These include the D1/H198Q [25], D1/T179H [26], and D2/I178H [28] mutants; FRL-PSII from *C. thermalis*, containing 4 Chl-*f* and 1 Chl-*d* [20]; and PSII from *A. marina*, containing 1 Chl-*a* (PD1) and 34 Chl-*d* [29].

Panel A in Fig. 5 presents the Tyr_Z_^•^Q_A_^•-^-*minus*-Tyr_Z_^•^Q_A_ difference spectra for PsbA3-PSII (black spectrum), D1/H198Q-PSII (red spectrum), D1/T179H-PSII (blue spectrum), and FRL-PSII from *C. thermalis* (green spectrum). As expected, the most significant spectral modification was observed in the D1/H198Q mutant, where the axial ligand of P_D1_ has been altered. In this mutant, the trough was shifted ∼3–4 nm to the blue, from 433–434 nm in PsbA3-PSII to 430 nm.

**Figure 5:**
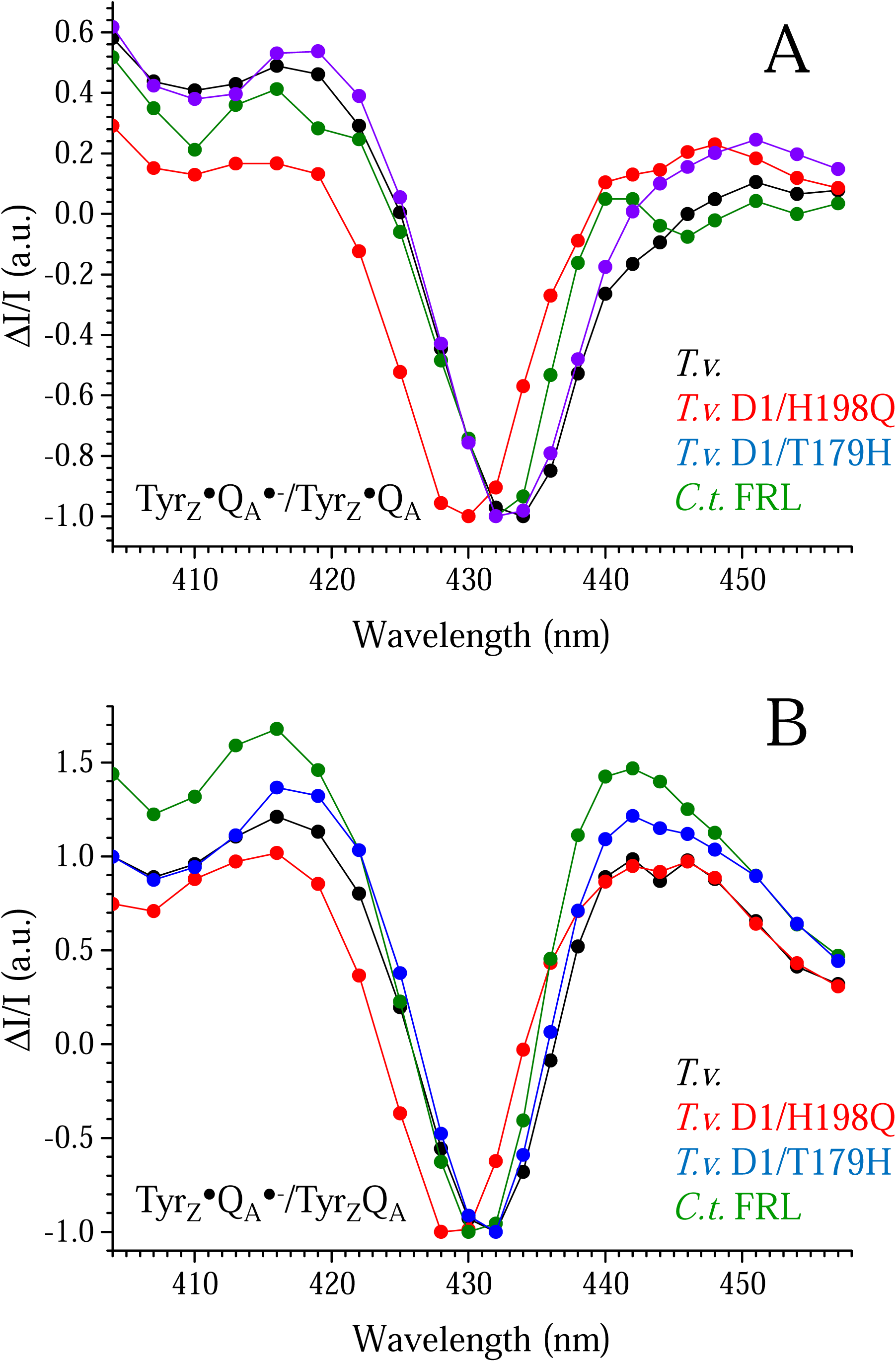
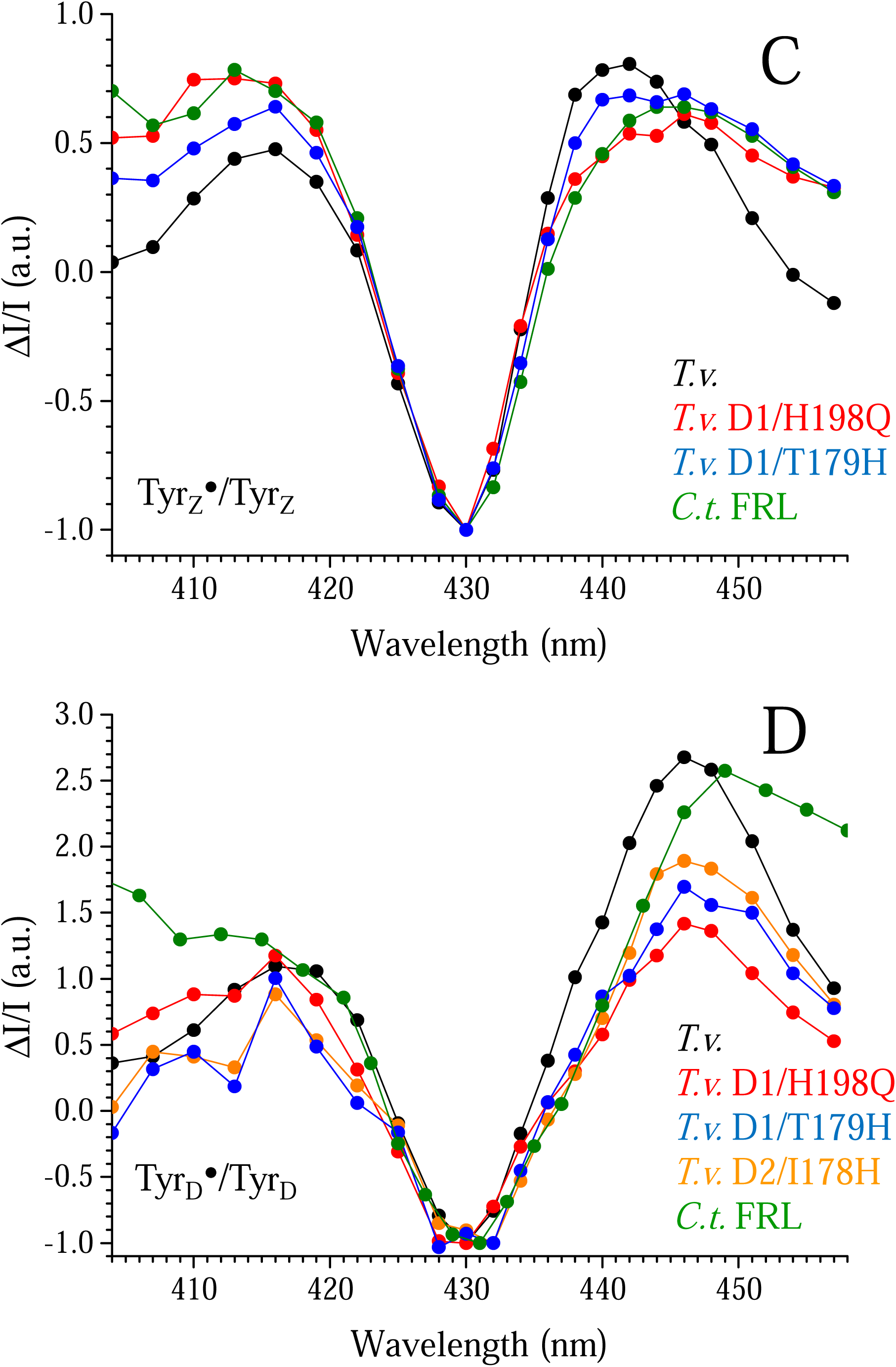
Panel A, spectra of the Tyr_Z_^•^Q_A_^•-^-*minus* -Tyr_Z_^•^Q_A_ difference in various PSII samples. Same conditions as in Panels A and B in Fig. 1. Panel B, spectra of the Tyr_Z_^•^Q_A_^•-^-*minus*-Tyr_Z_Q_A_ difference in various PSII samples. Same conditions as in Panels A and B in Fig. 1. Panel C, spectra of the Tyr_Z_^•^- *minus*-Tyr_Z_ difference in various PSII samples. Same conditions as in Panels A and B in Fig. 1. Panel D, spectra of the Tyr_D_^•^- *minus* -Tyr_D_ difference in various PSII samples. Same conditions as in Panels A and B in Fig. 1. All the spectra were normalized to -1 for the amplitude of the trough.

The Tyr_Z_^•^Q_A_^•-^-*minus*-Tyr_Z_^•^Q_A_ difference spectrum in PsbA3-PSII displayed a shoulder at 443–444 nm. This shoulder was not shifted in the D1/H198Q-PSII. However, in FRL-PSII, where Chl_D1_ is likely a Chl-*d*, this shoulder was shifted ∼3 nm to the red. This shift was accompanied by a narrowing of the spectrum between 434 nm and 440 nm, confirming the red shift of a spectral component. In the D1/T179H mutant, where Chl_D1_ is also affected, the shoulder at 443–444 nm also appeared modified, albeit less prominently.

Panel B in Fig. 5 displays the Tyr_Z_^•^Q_A_^•-^-*minus*-Tyr_Z_Q_A_ difference spectra for PsbA3-PSII (black spectrum), D1/H198Q-PSII (red spectrum), D1/T179H-PSII (blue spectrum), and FRL-PSII from *C. thermalis* (green spectrum). The shape of these spectra suggests that two pigments absorbing near 430 nm are affected by the formation of Tyr_Z_^•^Q_A_^-^, with one pigment exhibiting a blue shift and the other a red shift. The effect of the D1/H198Q mutation indicates that one of these two pigments is clearly P_D1_. In a previous study conducted on O - evolving PSII at pH 6.5 [41], we interpreted similar spectra associated with Tyr_Z_^•^Q_A_^•-^ formation as arising from a red shift in the Soret band of P_D1_ and a blue shift in the Soret band of Chl_D1_. Unfortunately, the signal-to-noise ratio of the current spectra, compared with those in [41], is insufficient to confirm or refute the involvement of Chl_D1_ in the Tyr_Z_^•^Q_A_^•-^-*minus* - Tyr_Z_Q_A_ difference spectrum under these experimental conditions.

Panel C in Fig. 5 shows the Tyr_Z_^•^- *minus* -Tyr_Z_ difference spectra for PsbA3-PSII (black spectrum), D1/H198Q-PSII (red spectrum), D1/T179H-PSII (blue spectrum), and FRL-PSII from *C. thermalis* (green spectrum). These spectra similarly suggest that more than one pigment absorbing around 430 nm is affected by the formation of Tyr_Z_^•^ compared to Tyr_Z_. Surprisingly, the spectrum remained unchanged in the D1/H198Q mutant, strongly suggesting that P_D1_ does not undergo an absorption shift upon Tyr_Z_^•^ formation.

This finding indicates that pigments other than P_D1_ experience electrochromic shifts in the presence of Tyr_Z_^•^. Despite averaging numerous spectra, the signal-to-noise ratio remained insufficient to unambiguously identify these pigments. However, given the spectral range in which the band shifts occur and the lack of effects in the mutants studied, it is plausible that a carotenoid or a chlorophyll other than P_D1_, Chl_D1_, or Chl_D2_ may be involved.

Panel D in Fig. 5 presents the Tyr_D_^•^-*minus*-Tyr_D_ difference spectra for PsbA3-PSII (black spectrum), D1/H198Q-PSII (red spectrum), D1/T179H-PSII (blue spectrum), D2/I178H-PSII (orange spectrum) [28], and FRL-PSII from *C. thermalis* (green spectrum). These spectra differ notably from the Tyr_Z_^•^- *minus*-Tyr_Z_ difference spectra. In particular, the Tyr_D_^•^-*minus*-Tyr_D_ difference spectra appear to involve only one pigment, with a clear shift from approximately 430 nm in the presence of Tyr_D_ to ∼446 nm in the presence of Tyr_D_^•^. The absence of changes in the difference spectra across the mutants tested suggests that neither P_D1_, Chl_D1_, nor Chl_D2_ is involved in this shift.

The situation in FRL-PSII (green spectrum) warrants further commentary. In dark-adapted FRL-PSII, the proportion of centers with a reduced Tyr_D_ is highly variable, with virtually no reduction observed in some cases. This variability was confirmed by the amplitude of the Tyr_D_^•^ signal in EPR measurements taken before and after illumination (data not shown). This suggests a significant difference compared to other PSII. While the underlying cause of this variability was not further investigated, it implies that the Tyr_D_^•^- *minus*-Tyr_D_ difference spectrum in FRL-PSII may be contaminated by a contribution from the Tyr_Z_^•^- *minus*-Tyr_Z_ difference spectrum.

Despite this, the positive part of the spectrum in the presence of Tyr_D_^•^ in FRL-PSII appears to shift to ∼449 nm, compared to ∼446 nm in the other samples. However, the fact that the pigment’s spectrum peaks at 430 nm in the presence of Tyr_D_, as observed in other PSII, indicates that the pigment itself does not have an altered spectrum. Instead, this suggests that the electrochromic effect is different in FRL-PSII. This difference likely arises from a modified electrostatic environment in FRL-PSII, as previously mentioned. Such modifications could also explain the distinct stability of Tyr_D_^•^ observed in these samples.

One pigment that has not been investigated until now in this report is P_D2_. To study it, we used PSII from *Synechocystis* 6803, in which the D2/H197A mutant has been constructed [30]. The results are shown in Fig. 6 for the four difference spectra. Surprisingly, even the Tyr_D_^•^-*minus*-Tyr_D_ difference spectrum remained unaffected by the D2/H197A mutation.

**Figure 6:**
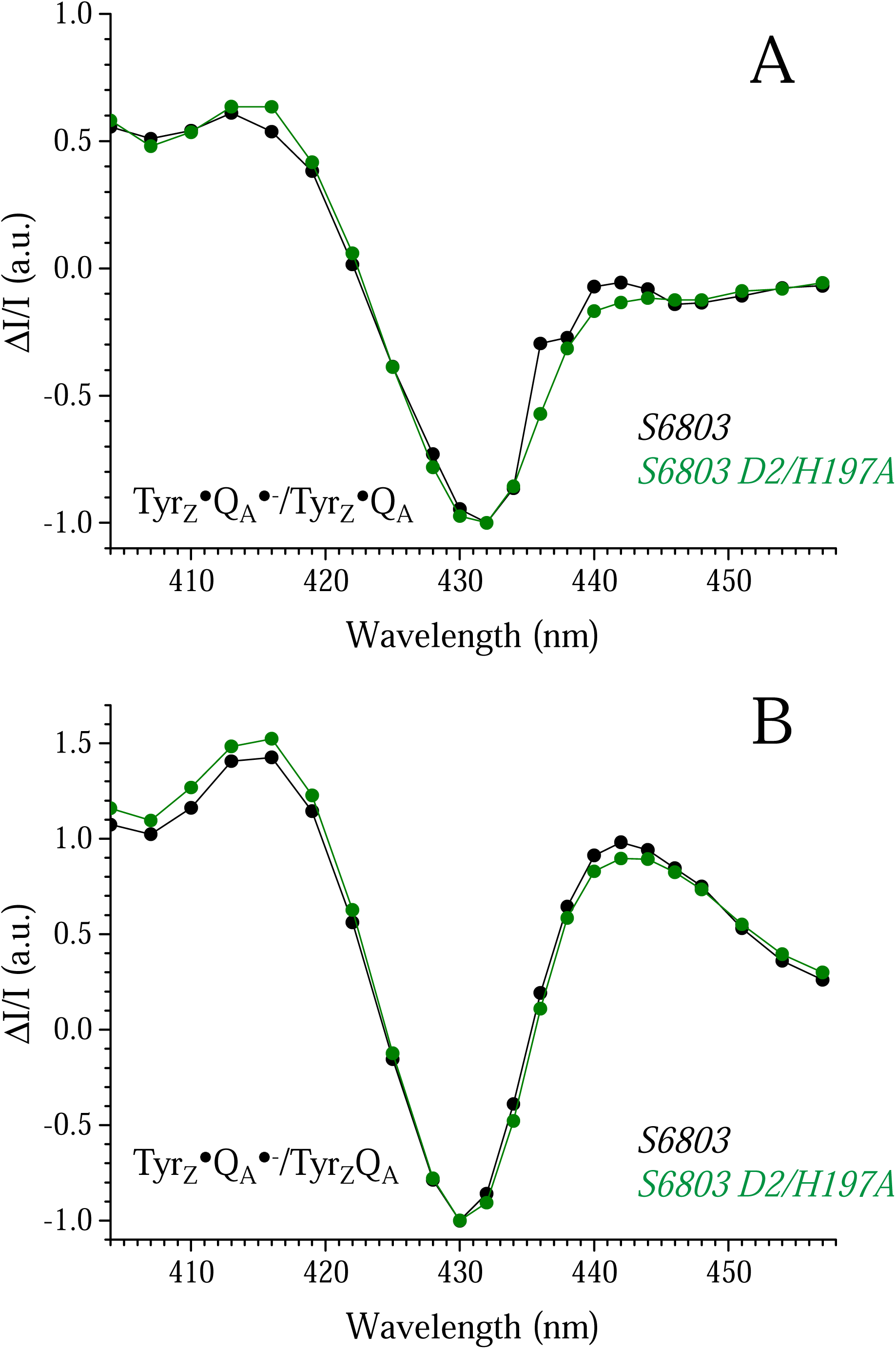
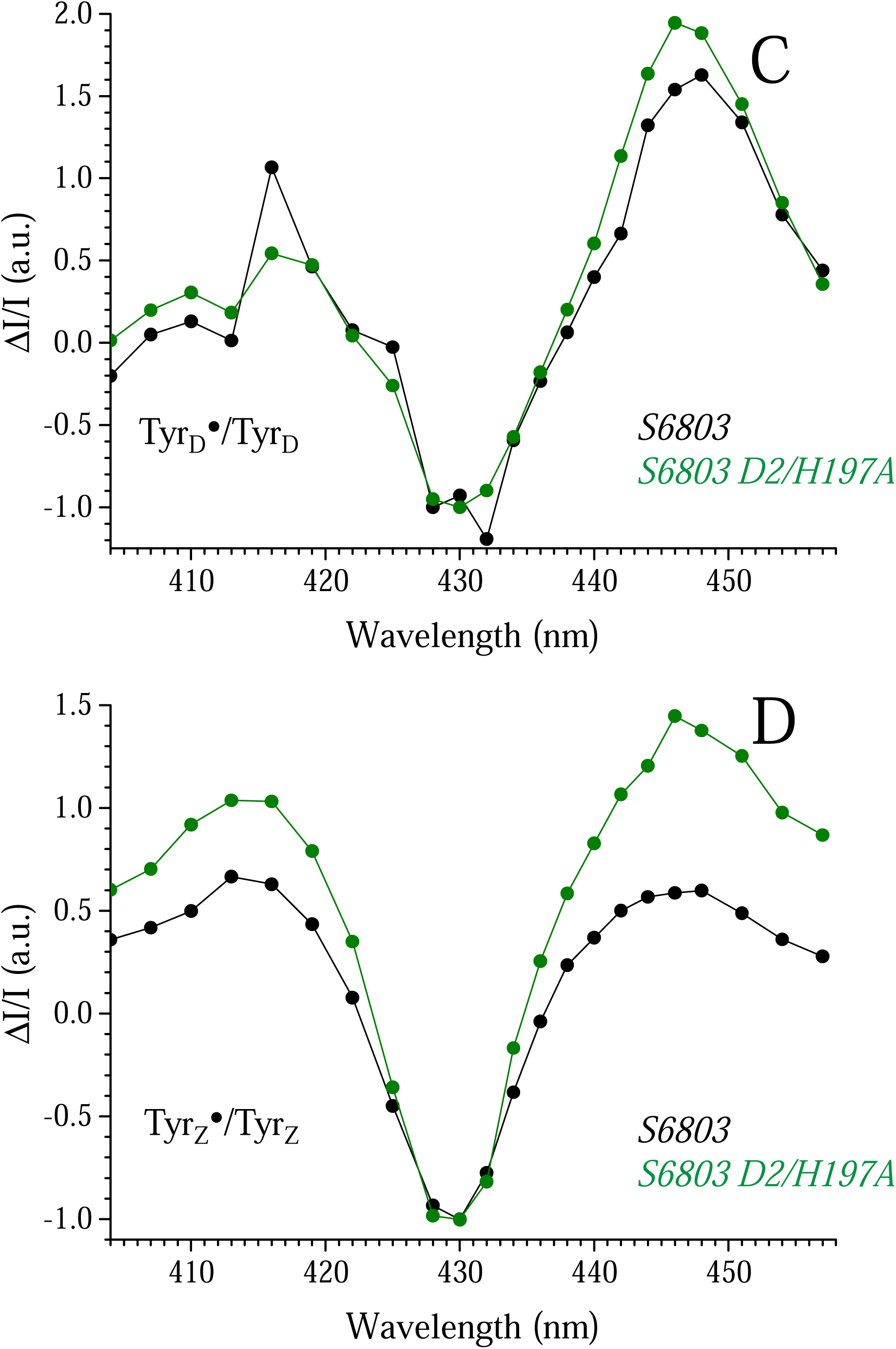
Panel A, spectra of the Tyr_Z_^•^Q_A_^•-^-*minus*-Tyr_Z_^•^Q_A_ difference in wild type *Synechocystis* 6803 (black spectrum) and in the D2/H197A mutant (green spectrum). Same conditions as in Panels A and B in Fig. 1. Panel B, spectra of the Tyr_Z_^•^Q_A_^•-^-*minus*-Tyr_Z_Q_A_ difference in wild type *Synechocystis* 6803 (black spectrum) and in the D2/H197A mutant (green spectrum). Same conditions as in Panels A and B in Fig. 1. Panel C, spectra of the Tyr_D_^•^-*minus*-Tyr_D_ difference in wild type *Synechocystis* 6803 (black spectrum) and in the D2/H197A mutant (green spectrum). Same conditions as in Panels A and B in Fig. 1. Panel D, spectra of the Tyr_Z_^•^-*minus*-Tyr_Z_ difference in wild type *Synechocystis* 6803 (black spectrum) and in the D2/H197A mutant (green spectrum). Same conditions as in Panels A and B in Fig. 1. All the spectra were normalized to -1 for the amplitude of the trough.

The lack of significant effects, except for the P_D1_ mutant D1/H198Q, in the two cases corresponding to the Tyr_Z_^•^Q_A_^•-^-*minus*-Tyr_Z_Q_A_ and Tyr_Z_^•^Q_A_^•-^- *minus* -Tyr_Z_^•^Q_A_ difference spectra suggests that, as mentioned above, either pigments other than P_D2_, Ch_D1_, and Chl_D2_ are affected, or the spectral changes are too weak given the signal-to-noise ratio of the experiments. One way to confirm this hypothesis is to investigate what happens in the PSII of *A. marina* with 34 Chl-*d* molecules and one Chl-*a*.

It was reported that the P_680_^•+^Q_A_^•-^-*minus*-P_680_Q_A_ difference spectrum at 77 K was identical to that in *T. elongatus* [31], which led the authors to confirm that P_D1_ was the Chl-*a*. Here, we examined the Tyr_D_^•^-*minu*s-Tyr_D_ and Tyr_Z_^•^- *minus* -Tyr_Z_ difference spectra. In *A. marina*, unlike in *T. vestitus*, Tyr_D_^•^ is not stable in the dark at pH 6.5, which led us to perform the difference spectra at pH 6.5. Of course, it would have been preferable to record them at pH 8.6, but, unfortunately, for a number of reasons beyond our control, we could not do so.

Fig. 7 shows the Tyr_D_^•^- *minus*-Tyr_D_ (Panel A) and Tyr_Z_^•^- *minus*-Tyr_Z_ (Panel B) difference spectra at pH 6.5, in FRL-PSII from *C. thermalis* (black spectra) and in PSII from *A. marina* (red spectra). As expected, Tyr_D_^•^ appeared stable at pH 6.5 in *C. thermalis*, with a Tyr_D_^•^-*minus*-Tyr_D_ difference spectrum completely flat in this PSII at this pH. In contrast, in *A. marina*, there was a large Tyr_D_^•^-*minus*-Tyr_D_ difference spectrum. One species whose spectrum is modified by the oxidation of Tyr_D_ absorbed at ∼455 nm. This supports the hypothesis that this species is a Chl-*d*. It is tempting to propose that this Chl-*d* has an absorption shifted to ∼470 nm in the presence of Tyr_D_^•^. However, the origin of the positive feature at ∼422 nm is less clear. For the Tyr_Z_^•^- *minus*-Tyr_Z_ difference spectra, the situation is even less clear-cut. While the possible presence of two spectral shifts in *A. marina* may be due to the doubling of the Soret band on either side of the Soret band for Chl-*a*, the differences between pH 6.5 and pH 8.6 in the case of *C. thermalis* remain to be understood.

**Figure 7:**
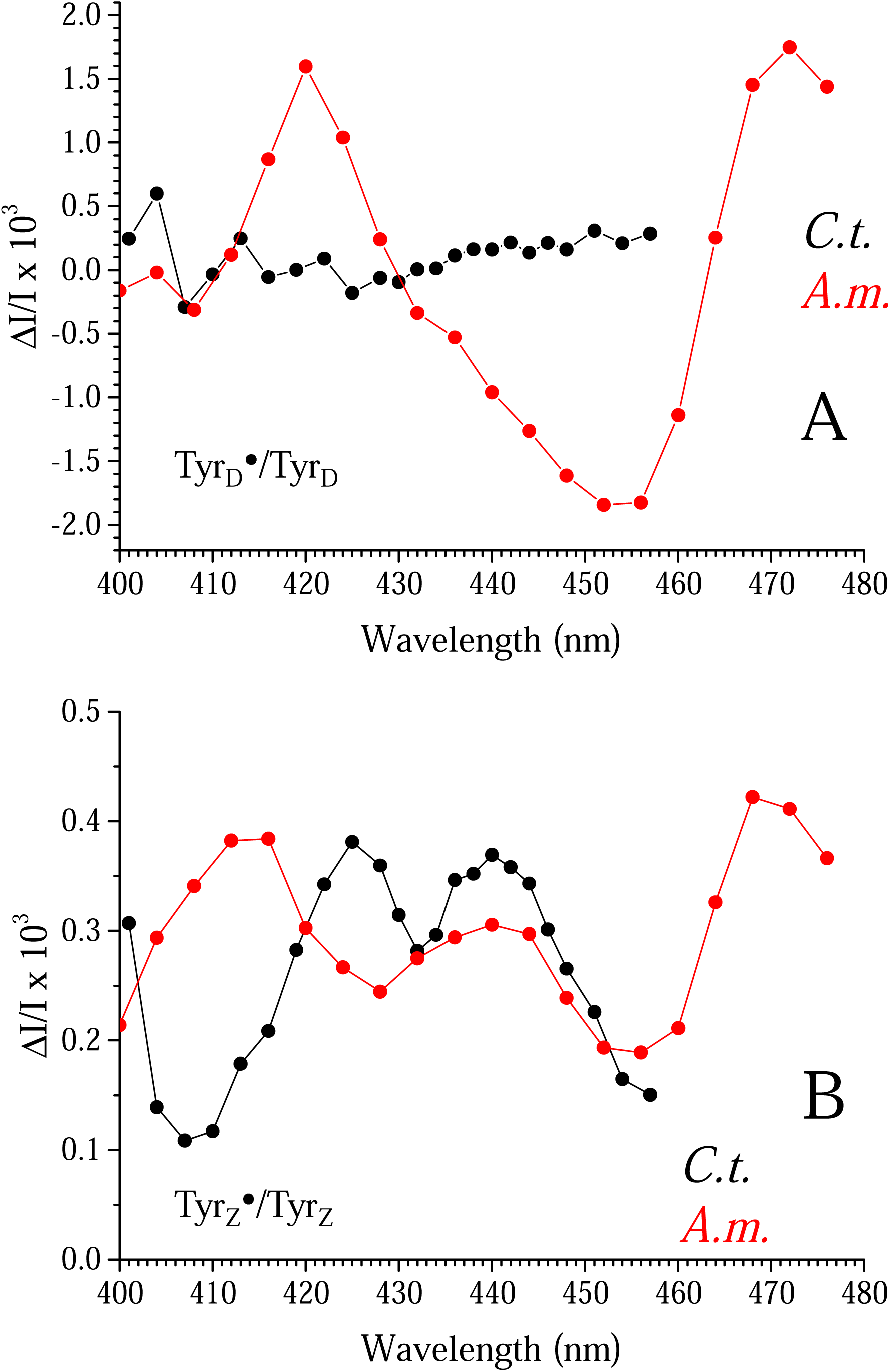
Panel A, spectra of the Tyr_D_^•^-*minus*-Tyr_D_ difference at pH 6.5 in *C. thermalis* (black spectrum) and in PSII from *A. marina* (red spectrum). Other conditions same as in Panels A and B in Fig. 1. Panel B, spectra of the Tyr_Z_^•^-*minus*-Tyr_Z_ difference at pH 6.5 in FRL-PSII from *C thermalis* (black spectrum) and in PSII from *A. marina* (red spectrum). Other conditions same as in Panels A and B in Fig. 1. The ΔI/I were averaged from the 6^th^ to 10^th^ flash.

Panel B in Fig. 1 shows that between 10 ms and 500 ms, a change in the ΔI/I occurred in the opposite direction when compared to the decay of Tyr_Z_^•^. Panel A in Fig. 8 shows an average of the ΔI/I from the 6th to the 10th flash in PsbA3-PSII from *T. vestitus* (black points), D1/H198Q-PSII (red points), and FRL-PSII from *C. thermalis* (green points). Panel B in Fig. 8 shows the spectra for the three PSII preparations. The three spectra are identical to the difference spectra Tyr_Z_^•^Q_A_^•-^- *minus* -Tyr_Z_Q_A_ head-to-tail. This suggests that the electrostatic environment resulting from the formation of Tyr_Z_^•^Q_A_^•-^ occurs with kinetics slower than the formation of the state. The different kinetics in Panel A of Fig. 8 could represent another manifestation of a different rearrangement of the electrostatic environment in the species studied here.

**Figure 8:**
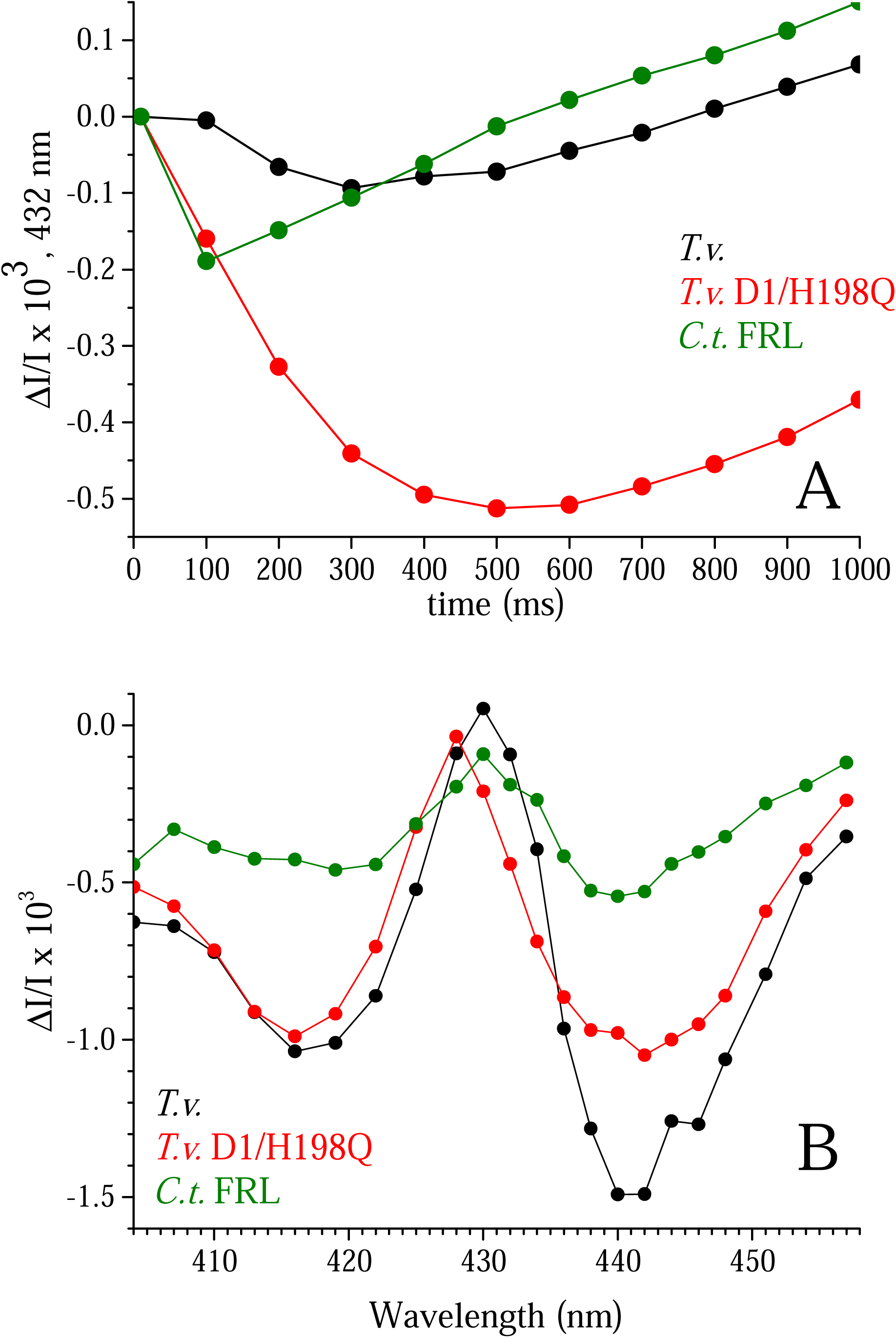
Panel A shows ΔI/I recorded at 432 nm and at pH 8.6 in Mn-depleted PSII from *C. thermalis* (green data points), from *T. vestitus* (black data points), and from the D1/H198Q mutant in *T. vestitus* (red data points). The ΔI/I values were averaged from the 6th to the 10th flash. The first point corresponds to the detector flash given 10 ms after the actinic flash. Panel B shows the wavelength dependence of the maximum amplitude of the kinetics from Panel A, measured at 300 ms for PsbA3-PSII from *T. vestitus*, at 100 ms for FRL-PSII from *C. thermalis*, and at 500 ms for D1/H198Q PSII from *T. vestitus*.

## Conclusion

The effects of Tyr_D_^•^, Tyr_Z_^•^, and Q_A_^•-^ radical formation on the absorption spectrum in the Soret region were studied in Mn-depleted Photosystem II (PSII) at pH 8.6. Flash-induced difference spectra were recorded for several PSII samples from: *i*) *Thermosynechococcus vestitus* (formerly *T. elongatus*), *ii*) *Synechocystis* sp. PCC 6803, *iii*) *Chroococcidiopsis thermalis* PCC7203 grown under far-red light, and *iv*) *Acaryochloris marina*. While P_D1_ is clearly the species whose spectrum is blue-shifted by ∼3 nm in the presence of Q_A_^•-^, the spectra recorded from these different samples suggest that P_D1_, P_D2_, Chl_D1_, Phe_D1_, and Chl_D2_ are not the species whose absorption spectrum is modified by the formation of Tyr_D_^•^ and Tyr_Z_^•^. The conclusion for Phe_D1_ is drawn from the similarity of the Tyr_Z_^•^- *minus*-Tyr_Z_ difference spectrum in PsbA3-PSII with E-130 and PsbA1/Y160F-PSII with Q-130, as shown in [42]. It is therefore proposed that other pigments, such as antenna Chl and/or Car near the reaction center, are involved. Additionally, it is shown that: *i*) there is no proton release into the bulk upon the oxidation of Tyr_D_ at pH 8.6, and *ii*) the rearrangement of the electrostatic environment of the pigments involved in the light-induced difference spectra in the samples studied, upon the formation of Tyr_D_^•^, Tyr_Z_^•^, and Q_A_^•-^, likely occurs differently from both kinetic and structural perspectives.

